# Rhizospheric miRNAs affect the plant microbiota

**DOI:** 10.1101/2022.07.26.501597

**Authors:** Harriet Middleton, Jessica Ann Dozois, Cécile Monard, Virginie Daburon, Emmanuel Clostres, Julien Tremblay, Jean-Philippe Combier, Étienne Yergeau, Abdelhak El Amrani

## Abstract

Recently, small RNAs have been shown to play important roles in cross-kingdom communication, notably in plant-pathogen relationships. Plant miRNAs were even shown to regulate gene expression in the gut microbiota. But what impact do they have on the plant microbiota? Here we hypothesized that plant miRNAs can be found in the rhizosphere of plants, where they are taken up by rhizosphere bacteria, influencing their gene expression, thereby shaping the rhizosphere bacterial community. We found plant miRNAs in the rhizosphere of *Arabidopsis thaliana* and *Brachypodium distachyon*. These plant miRNAs were also found in rhizosphere bacteria, and fluorescent synthetic miRNAs were taken up by cultivated bacteria. A mixture of five plant miRNAs modulated the expression of more than a hundred genes in *Variovorax paradoxus*, whereas no effect was observed in *Bacillus mycoides*. Similarly, when *V. paradoxus* was grown in the rhizosphere of *Arabidopsis* that overexpressed a miRNA, it changed its gene expression profile. The rhizosphere bacterial communities of *Arabidopsis* mutants that were impaired in their miRNA or small RNA pathways differed from wildtype plants. Similarly, bacterial communities of *Arabidopsis* overexpressing specific miRNAs diverged from control plants. Finally, the growth and the abundance of specific ASVs of a simplified soil community were affected by exposure to a mixture of synthetic plant miRNAs. Taken together, our results support a paradigm shift in plant-bacteria interactions in the rhizosphere, adding miRNAs to the plant tools shaping microbial assembly.

## Introduction

MicroRNAs (miRNAs) are small ∼21nt non-coding RNAs that control target gene expression, through sequence complementarity. Their roles in plants vary from regulating developmental processes to responding to abiotic and biotic stresses. Plants also use miRNAs to interact with pathogens: for example, cotton plants use them to inhibit the fungal pathogen *Verticillium dahlia* [1] and *Arabidopsis thaliana* and tomato to inhibit another fungal pathogen, *Botrytis cinerea* [2–4]. The more general role of plant miRNAs in shaping the microbiota is, however, not known. We had hypothesized that miRNAs could be a novel, key mediator in the rhizosphere [5], as it was shown to be in the mammalian gut. In mice and human, both intestinal miRNAs and ingested plant-derived miRNAs regulate gut bacterial gene expression and growth, shaping the gut microbiota [6, 7] [8].

*Arabidopsis* roots contain many miRNAs, of which over half are expressed in a tissue-specific manner, with several being enriched at the root tip, in the early meristematic zone [9]. This zone is also a recognized hotspot for plant-driven microbial selection [10]. The current paradigm is that the plant selects the rhizospheric microbiota through rhizodeposition, including exudates such as sugars, peptides, amino acids, nucleic acids, nucleotides, fatty acids or secondary metabolites [11]. Here, we hypothesize that plant miRNAs 1) can be found in the rhizosphere of plants, 2) are taken up by rhizosphere bacteria, 3) influence the gene expression of rhizosphere bacteria, and 4) shape the bacterial community. We report several independent experiments designed to test these hypotheses.

## Methods

*The full method description is available as supplementary material*.

### Detection of plant miRNAs in the rhizosphere and roots

For rhizosphere analyses, triplicate *Arabidopsis thaliana* (Col-0) and *Brachypodium distachyon* (Bd21-3) were grown for a month, alongside unplanted soils. For root analyses, we grew ten replicated *Aradidopsis thaliana* (Col-0) plants for 21 days, with unplanted controls. RNA was extracted from the plant roots and rhizosphere and from the unplanted soils and sent for small RNA sequencing. Sequences between 18 and 27 nucleotides were mapped against *Arabidopsis* and *Brachypodium* genomes and were assigned to known miRNAs. From these miRNAs we constructed an abundance table that was used to compare the rhizosphere to the unplanted controls. Rhizospheric and root miRNAs were defined as the miRNAs that had at least 10 reads for each of the rhizosphere or root samples and a maximum of one read across all the bulk soil samples.

### Internalization of plant miRNAs by bacteria

#### Detection of plant miRNAs in rhizospheric bacteria

We extracted bacterial cells from the rhizosphere soil of one-month old *A. thaliana* (Col-0) using a Nycodenz gradient. RNA was extracted from the bacterial pellet [12], sequenced and processed as described above, but with a lower threshold because of the lower number of plant miRNA sequences retrieved. Only the miRNAs that were represented by at least 5 reads in each rhizosphere samples and absent in the bulk soil samples were kept. To further confirm that miRNAs found in the rhizospheric bacteria originated from the plant and not the bacteria, miRNA sequences were searched for on the + and - strands of 3,837 bacterial genomes [13], of which 1,160 were isolated from plants.

#### Confocal microscopy

*Variovorax paradoxus* EPS and *Bacillus mycoides* YL123 were grown overnight in liquid media. The cultures were incubated for 4 hours with 3’-Cy5 fluorescent miRNAs (ath-miR159a or a scrambled control – same nucleic acid content but in different order), or a pCp-Cy5 control. Twenty minutes before visualization, the cultures were also treated with MitoTracker Green FM (Invitrogen). We visualized the washed and concentrated culture using a confocal microscope (Zeiss LSM780).

#### Flow cytometry

We prepared the cultures as described in the confocal microscopy section. The cells were fixed with 4% Paraformaldehyde (PFA) and then stained with a DNA marker (Hoechst 33342). The cells were diluted in PBS and processed by flow cytometry (BD LSRFortessa). We used a variety of controls to ensure that our statistical analyses were conducted on bacteria that were positive for our green (MitoTraker), red (Cy5 tagged miRNAs) and blue (Hoechst) markers.

### Effect of miRNAs on bacterial gene expression

#### Bacterial transcriptomic response to miRNAs

*Variovorax paradoxus* EPS and *Bacillus mycoides* YL123 were grown until the exponential phase, when they were exposed to a mix of the 6 most abundant rhizospheric miRNAs: miR159a, miR159b, miR159c, miR161.1, miR158a and miR165b, or a control mix of 6 scrambled miRNAs. The cultures were sampled after 20 min and 120 min of incubation, pelleted, and the RNA of the cell pellet was extracted and sequenced. The transcripts were mapped on their respective genomes, and differential expression analyses (plant miRNA vs. scrambled miRNA) were done using DESeq2.

#### In silico analysis of miRNA targets

To predict potential targets of the six miRNA used on the genome of *Variovorax paradoxus*, we implemented a workflow [14] named *mirnatarget 1.0 : miRNA target finder* (https://github.com/jtremblay/MiRNATarget), which was inspired from the plant miRNA target finder, *psRNAtarget* [15].

#### In vitro miPEP treatment and transcriptomic experiment

Approximately 50 *Arabidopsis thaliana* (Col-0) surface-sterilized seeds were grown axenically in Petri dishes in a growth chamber. After twenty days of growth, the plants were treated twice within 24 hours by inoculating miPEP159c, a scrambled miPEP, or water at the plant crown. One hour after the second miPEP treatment, *Variovorax paradoxus* was inoculated along the roots of the seedlings. Two hours after the bacterial inoculation, the plants and rhizospheres were sampled, and RNA was extracted. From the potential targets of miR159c found above, three were selected for RT-qPCR analyses in the rhizosphere – alpha-2-macroglubulin, phosphatidate cytydylyltransferase, and LysR. We also quantified the abundance in plant tissues of the primary transcript to mir159c, pri-miR159c.

### Effect of miRNAs on the bacterial community

#### Arabidopsis mutant experiment

Five *A. thaliana* mutants were chosen. *RTL1* mutant over-expresses RTL1 protein which results in a suppression of siRNA pathway without affecting miRNAs [16]. *RTL1myc* over-expresses RTL1 protein flagged with Myc epitope, rendering RTL1 less active, so siRNA pathway is less suppressed than with *RTL1* mutant. *Ago1-27* mutant has AGO protein function partially impaired and is completely post-transcription gene silencing (PTGS) deficient [17]. *Dcl1-2* mutant has total loss of function of DCL1 protein resulting in low levels of miRNA and developmental problems [18]. *Hen1-4* mutant is miRNA defective but is also affected in some siRNA – PTGS [19]. HEN1 methylates siRNA and miRNA to maintain their levels and size, but also to protect them from uridylation and subsequent degradation [20]. The plants were grown for a month, after which the roots and attached rhizosphere were sampled, its DNA extracted, amplified using 16S rRNA gene primers, and sequenced. The same primers were used in real-time quantitative PCR to quantify bacterial abundance. Amplicon sequencing data was processed with AmpliconTagger [21] and the R package “phyloseq” v 1.32.0 [22].

#### miPEP experiment

We treated *Arabidopsis thaliana* Col-0 with 500 µL of water (control condition) or a miPEP solution (20 µM of miPEP159a, miPEP159b, or miPEP159c), applied at the base of the crown, 3 times a week for a total of 10 applications. We then extracted the DNA from the roots and the attached rhizosphere and sequenced and quantified the 16S RNA gene as described above.

#### Simplified soil community experiment

We created a simplified soil community by inoculating five different growth media with 2 g of agricultural soil. The cultures were normalized to the same optical density, pooled, pelleted and suspended in PBS. The cells were inoculated in a 96-wells plate containing a mixture of 17 amino acids as nitrogen source. Five wells were treated with a mixture of rhizospheric miRNAs (ath-miR158a-3p, ath-miR158b, ath-miR159a, ath-miR827, and ath-miR5642b) and five wells were treated with a mixture of scrambled miRNAs. These miRNAs were all found in the rhizosphere (but for some below the stringent threshold used above) and were predicted to target bacterial genes associated with nitrogen cycling. We measured bacterial growth every hour for 52 hours, after which we sampled the bacteria, extracted the DNA, and amplified and sequenced the 16S rRNA gene as described above.

## Results

### Plant miRNAs are present in the rhizosphere

We sequenced small RNA extracted from *A. thaliana* rhizosphere soil and unplanted soils. One-hundred-eleven ath-miRNAs (mapped on *A. thaliana*’s genome) were detected, of which 14 were present in the rhizosphere with more than 10 reads per sample and absent in unplanted soil (Fig. 1a). The most abundant miRNAs in the rhizosphere were ath-miR158a-3p, ath-miR161.1, and various members of the miR159, miR166 and miR165 families (Fig. 1a). We then sequenced the rhizosphere miRNAs of a second model plant, *Brachypodium distachyon*. Ten bdi-miRNAs (mapped on *B. distachyon*’s genome) out of 81 miRNAs were present in the rhizosphere with more than 10 reads per sample and absent in unplanted soil (Fig. 1b). The most abundant miRNAs were bdi-miR159b-3p, bdi-miR156, bdi-miR166, bdi-miR396, and bdi-miR167. Amongst the rhizospheric miRNAs detected, four were common between *A. thaliana* and *B. distachyon*: miR159b-3p, miR167, miR166, and miR396.

**Figure 1:**
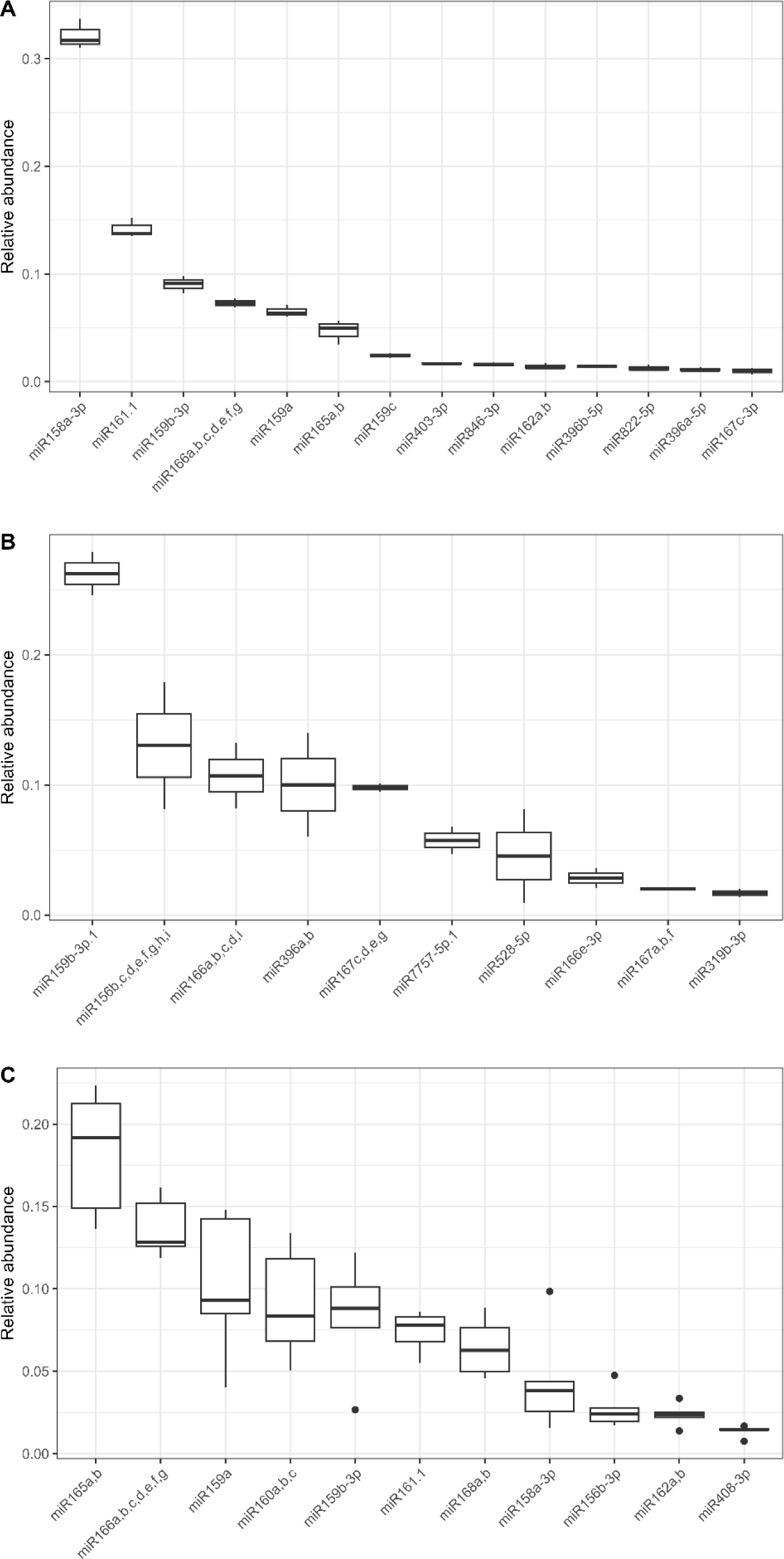
Plant miRNAs are present in the rhizosphere and absent in unplanted soil. Relative abundance of the plant miRNAs found in the rhizosphere of *Arabidopsis thaliana* (A), *Brachypodium distachyon* (B) and in the roots of *Arabidopsis thaliana* (C), while completely absent from unplanted soil.

We then grew more *Arabidopsis* plants and sequenced their root miRNAs to confirm that the rhizospheric miRNAs could be coming from the plant roots. Eleven miRNAs were represented by at least 10 reads in each root sample. There was a clear dominance of ath-miR165, ath-miR166, ath-miR159a, ath-miR160, and ath-miR159b.3p (Fig. 1c). Among these 11 miRNAs, 7 were common with the miRNAs found in *Arabidopsis* rhizosphere (ath-miR158a-3p, ath-miR159a, ath-miR159b-3p, ath-miR161.1, ath-miR162, ath-miR165, ath-miR166), which included most of the top five most abundant root and rhizosphere miRNAs.

### *A. thaliana* miRNAs are internalized by soil bacteria

Bacterial cells were isolated from the rhizosphere of one month-old *A. thaliana*, washed and their RNA content was extracted and sequenced to identify potentially internalized plant miRNAs. The miRNAs were mapped against *A. thaliana* TAIR10.1 genome, identifying a total of 34 ath-miRNAs. Five miRNAs – namely ath-miR158a-3p, ath-miR161.1, miR162, ath-miR159b-3p and ath-miR159a – were represented by at least 5 reads per rhizosphere samples and absent in bacteria extracted from the unplanted soil (Fig. 2a). Four out of these five miRNAs were among the five most abundant miRNAs found in the rhizosphere of *A. thaliana* (Fig. 1a) and were found in a similar order of relative abundance. To ensure that these miRNAs did not come from the bacteria themselves, their sequences were searched for in 3,837 soil bacterial genomes, of which 1,160 were of bacteria isolated from plants [13]. No matches were found, meaning that the miRNAs detected in the bacteria could not be produced by the bacteria.

**Figure 2:**
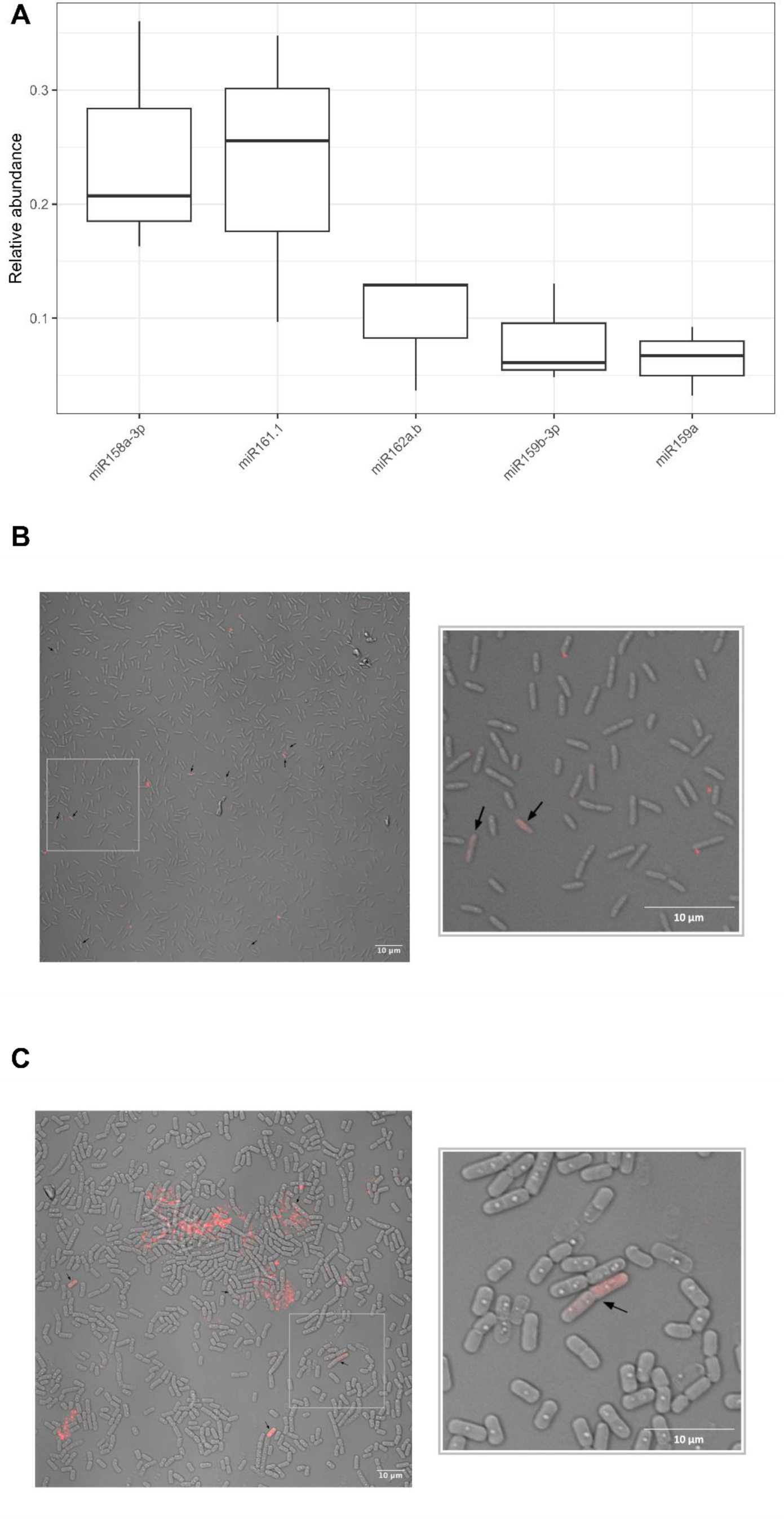
Plant miRNAs are internalized by rhizosphere bacteria. Relative abundance of *Arabidopsis thaliana* miRNAs found in cells pelleted from the rhizosphere and absent in cells pelleted from unplanted soil (A) and confocal microscopy images of *Variovorax paradoxus* (B) and *Bacillus mycoides* (C) after a 4-hour exposure to ath-miR159a tagged with Cy5. Arrows indicate cells containing the fluorescent molecule.

We then exposed two typical rhizosphere bacteria, *Variovorax paradoxus* EPS [23] and *Bacillus mycoides* [24] to a Cy5-tagged synthetic ath-miR159a and visualized its localization using confocal microscopy. Images show a clear localization of the miRNA inside many bacterial cells (Fig. 2b-c). Flow cytometry confirmed that an average of 6.51% *Variovorax* cells contained the Cy5 signal from plant miRNAs (average median fluorescence intensity= 123.00), compared to 4.95% of *Bacillus* cells (average median fluorescence intensity= 85.75). The scrambled miRNA, containing the same nucleotides as ath-miR159a but in a different order, was also internalized (Supplementary Figure S1), suggesting a general sequence-independent internalization mechanism for miRNAs. *Variovorax*, in contrast to *Bacillus*, internalized more efficiently the tagged miRNAs than the tagged single nucleotide. Indeed, a comparable amount of pCp-Cy5 was also internalized by *Bacillus* (on average 5.04%, average median fluorescence intensity=75.8), but this was an order of magnitude lower for *Variovorax* (on average 0.67%, average median fluorescence intensity=81.5) (Supplementary Figure S1 and S2). This suggests that *Variovorax* actively internalized the miRNAs, whereas it might have passively entered *Bacillus* either as intact miRNAs or degradation products of the miRNA.

### Plant miRNAs shifts rhizosphere bacterial gene expression

We then incubated *Variovorax* and *Bacillus* with a synthetic mixture of the six most abundant *A. thaliana* rhizosphere miRNAs: miR159a, miR159b, miR159c, miR161.1, miR158a and miR165b, or a mixture of scrambled miRNAs at the same concentration. *Bacillus* did not respond to the treatment – no gene was significantly differentially expressed following incubation with the synthetic miRNAs. In contrast, *Variovorax* showed important changes in response to the miRNA confrontation. After 20 min of incubation, the expression of 79 genes was significantly lower in the plant miRNA-treated cultures and the expression of 44 genes was significantly higher (adjusted *P*<0.05, Fig. 3a). After 120 min of incubation, the expression of 24 genes was significantly lower in the plant miRNA-treated cultures and the expression of 104 genes was significantly higher. Many genes were repressed after 20 min following the addition of the synthetic plant miRNAs to the bacterial culture, whereas after 120 min, more genes presented an increased than decreased expression. Only one gene was differentially expressed at both time points, a gene coding for a methionine synthase (VARPA_RS01000), which was overexpressed in response to the plant miRNA treatment.

**Figure 3:**
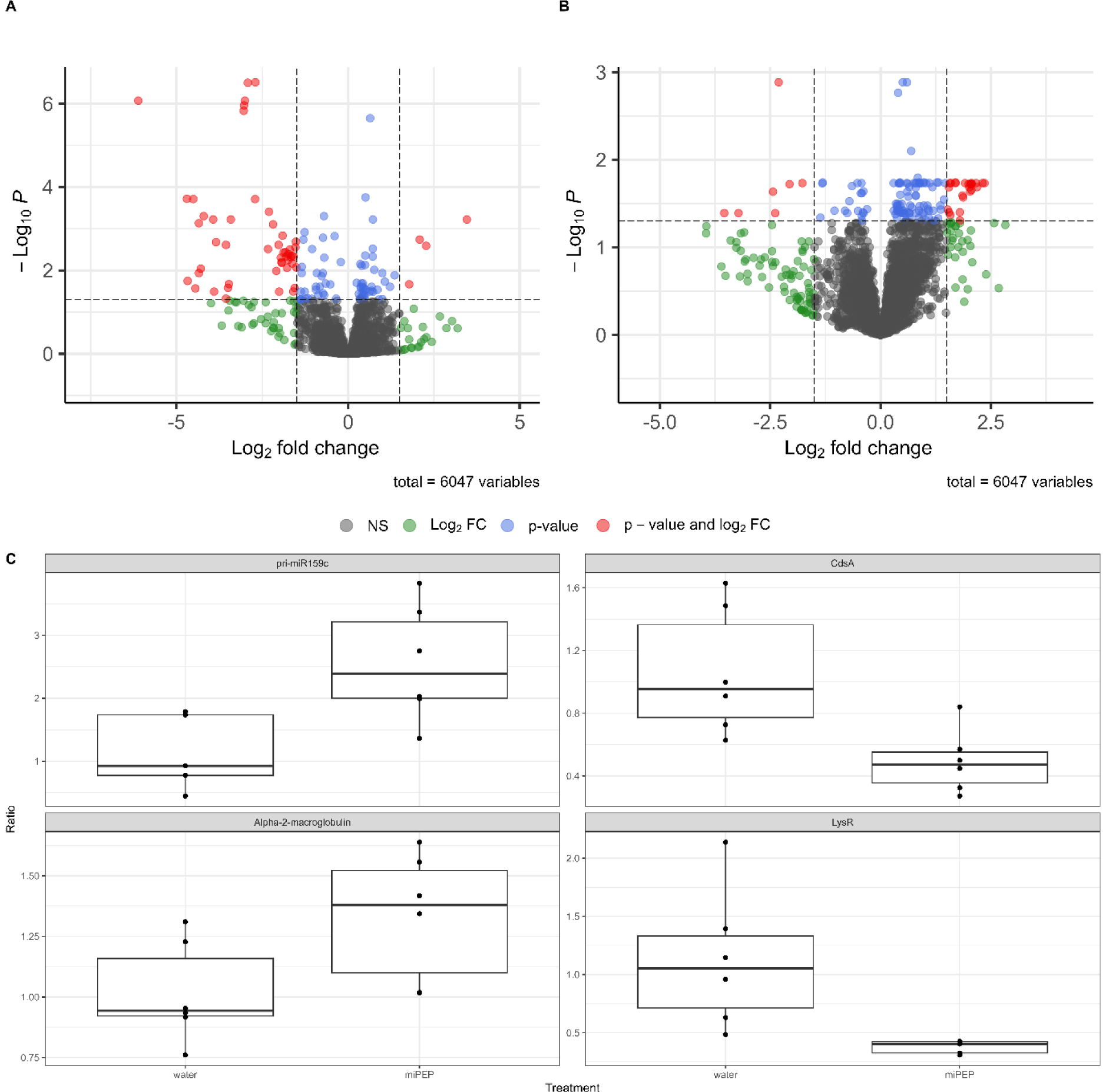
Plant miRNAs affect the transcriptome of a rhizosphere bacterium. Gene expression of *Variovorax praradoxus* after 20 min (A) and 120 min (B) exposure to a mixture of five synthetic miRNAs. (C): Arabidopsis thaliana expression of pri-miR159c and rhizospheric *Variovorax paradoxus* expression of CdsA, Alpha-2-macroglobulin and LysR after exposure of the plant to miPEP159c.

Using the rules of a plant small RNA target finder, psRNAtarget, we compared the differentially expressed genes from the previous experiment with the predicted target genes for the six miRNAs used. The six miRNAs were predicted to target 237 sequences in the *Variovorax paradoxus* EPS genome. Amongst these, 100 targets were positioned too far from any coding sequence (CDS), so they were removed from the following analysis, resulting in 137 potential targets. Amongst the 123 genes differentially expressed at 20 min, only two were predicted as targets *in silico*: VARPA_RS05960 (targeted by miR165), coding for a L-iditol 2-dehydrogenase, and VARPA_RS26385 (targeted by miR159a, b and c), coding for a phosphatidate cytidylyltransferase. Only two genes were predicted as targets amongst the 128 genes at 120 min: VARPA_RS00680 (targeted by miR165), coding for a hypothetical protein, and VARPA_RS22555 (targeted by miR158a-3p and miR159c), coding for a non-ribosomal peptide synthetase.

To confirm the effect of miRNAs on the bacterial transcriptome *in planta*, we inoculated the miRNA-encoded peptide (miPEP) miPEP159c to *Arabidopsis* plants growing *in vitro* and inoculated with *Variovorax*. miPEPs increase the expression of specific plant miRNAs [25, 26]. We selected miR159c because it was among the most abundant miRNAs in the rhizosphere, was in the mixture of miRNAs that modulated the gene expression of *Variovorax* and was predicted to target several key genes. The relative expression of the corresponding precursor miRNA (pri-miR159c) in the *Arabidopsis* plant tissue increased as compared to the water control (t-test: t=3.31, P= 0.0145; Fig. 3c). We then quantified the expression of three *Variovorax* genes determined to be potential targets of the miR159c according to our bioinformatic and transcriptomic analyses. One hundred and twenty minutes after the miPEP159c application, the relative expression of the LysR and the phosphatidate cytidylyltransferase (CdsA) genes decreased by a factor 2.69 and 2.16, respectively (t-test: t=4.27, *P*=0.00537, and t=3.43, P=0.00647, respectively, Fig. 3c), whereas the expression of alpha-2 macroglobulin gene increased by a factor 1.30 (t-test: t=2.92, P=0.0462, Fig. 3c) in comparison with the water control. The relative expressions of the three genes and the pri-miR159c following the application of the scrambled miPEP control (same amino acid composition as the miPEP, but in different order) were not significantly different from the water control.

### Plant miRNAs influence the rhizosphere bacterial community

To investigate the role of plant small RNAs on the rhizospheric microbial diversity, we grew *A. thaliana* mutants with disturbed miRNA and/or siRNA biosynthesis pathways and analyzed their rhizospheric microbial communities by 16S rRNA gene amplicons sequencing. The rhizosphere bacterial communities varied across the different genotypes (Permanova: *P*<0.05; Fig 4a). In PCoA ordinations, *ago1-27* and *RTL1myc* mutants’ communities were more like unplanted soil communities than those of WT plants (not shown). Bacterial diversity was higher in the rhizosphere of most mutant plants as compared with the WT plants (Fig. 4b).

**Figure 4:**
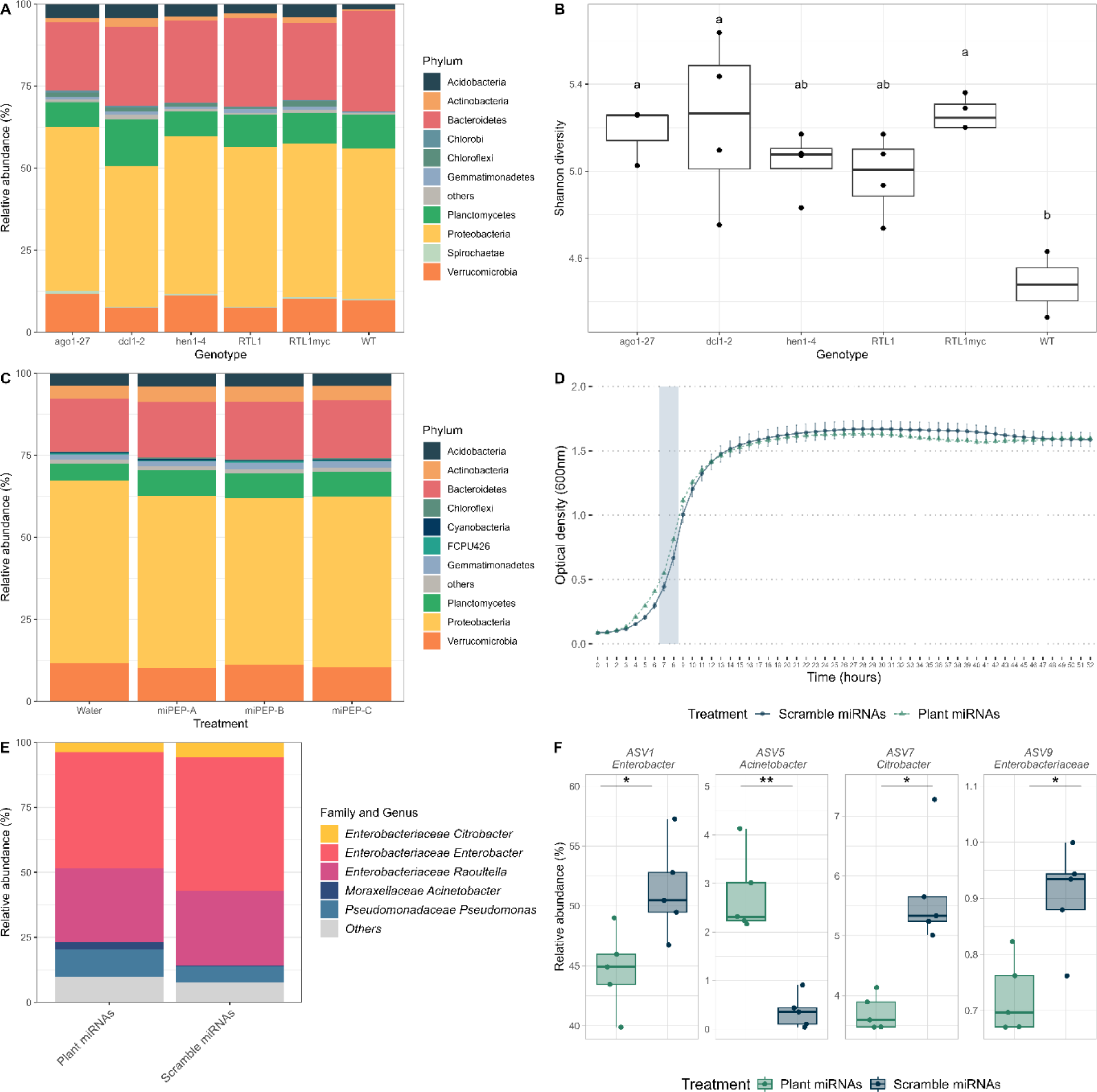
Plant miRNAs affect the bacterial community. Phylum-level bacterial community composition (A) and Shannon diversity (B) for different *Arabidopsis* mutants with impaired small RNAs pathways. (C): Phylum-level bacterial community composition for *Arabidopsis* plants inoculated with miPEP159a, miPEP159b or miPEP159c. Growth curve (D), community composition at genus level (E), and relative abundance of significantly affected ASVs (F) for a simplified soil microbial community exposed to a mixture of synthetic plant or scrambled miRNAs

We then treated soil-grown *Arabidopsis* with miPEPs to further test for the effect of the overexpression of specific miRNAs on the rhizospheric bacterial community. The application of three miPEPs (miPEP159a, miPEP159b, and miPEP159c) on the crown of *Arabidopsis* growing in soil changed the bacterial community in the roots/rhizoplane (Permanova: R^2^=0.160, P=0.031). All pairwise comparisons with the water control were significant or nearly significant (pairwise Permanova: miPEP159a R^2^=0.106, adjusted P=0.083; miPEP159b R^2^=0.164, adjusted P=0.025; miPEP159c R^2^=0.154, adjusted P=0.033). At the phylum level, the application of the miPEPs increased the relative abundance of the *Proteobacteria* (F=8.91, P=0.00029) and decreased the relative abundance of the *Planctomycetes* (F=40.78, P=3.64 x 10^−10^) (Fig. 4c). This was due to significant (adjusted P <0.05) differences between the water control and each of the miPEP treatments in post-hoc Tukey HSD tests. The application of the miPEP on the crown of *Arabidopsis* did not, however, affect the bacterial alpha diversity nor abundance.

Finally, we subjected *in vitro* a simplified microbial community that was enriched from an agricultural soil to a mix of five synthetic plant miRNAs (ath-miR158a-3p, ath-miR158b, ath-miR159a, ath-miR827, and ath-miR5642b), or a mix of their scrambled counterparts. Exposure to the plant miRNAs significantly disturbed the growth of the microbial communities during the log phase (t-test: P<0.05, Fig. 4d). The bacterial community contained twenty ASVs at the endpoint of the incubation (52 hours) and we found shifts in the bacterial community composition due to the plant miRNA exposure (Fig. 4e) and significant shifts in four ASVs related to the *Enterobacter*, *Acinetobacter*, *Citrobacter* genera, and to the *Enterobacteriaceae* family (Fig. 4f).

## Discussion

Using multiple lines of independent evidence, we confirmed our four hypotheses. Plant miRNAs are present in the rhizosphere (1) and are taken up by rhizobacteria (2), which induce changes in their transcriptome (3), leading to shifts in the bacterial community (4). Plants and pathogenic fungi interact using small RNAs [1–4, 27–29] and gut bacteria are influenced by host miRNAs [6, 7], but this is the first report of this mechanism for plant-bacterial community interactions in the rhizosphere.

We detected for the first time plant miRNAs in the rhizosphere. The two model plants, *A. thaliana* and *B. distachyon* harbored a similar complement of plant miRNAs in their rhizospheres. Although it would require confirmation from more plant species, the presence of similar miRNAs in the rhizosphere of a dicotyledon and a monocotyledon suggests a conserved feature among land plants. All the major miRNAs that we found in the rhizosphere of *Arabidopsis* were also detected in the roots. This agrees well with previous reports of root miRNAs, where the two most abundant rhizospheric miRNAs ath-miR158a and ath-miR161.1, were highly enriched in the early meristematic zone [9]. Many root exudates, such as extracellular DNA, soluble compounds, and mucilage, are produced and secreted, by border cells, in this region of the root tip [30]. Even though there was a large overlap between the root and rhizosphere miRNAs in our two experiments, the relative abundance of the miRNAs was not the same, alluding to a potential selection mechanism for the miRNAs that make it to the rhizosphere. Alternatively, this pattern could also be explained by different half-life in the rhizosphere, preferential uptake by bacteria, the sequencing of entire roots, or by the slightly different conditions under which the two experiments were run.

Bacteria growing in the rhizosphere of *Arabidopsis* contained plant miRNA inside their cells and isolated rhizosphere bacteria took up a fluorescent synthetic plant miRNA. The sequence of the miRNA did not affect the uptake, as the scrambled miRNA was taken up just as efficiently as the plant miRNA. Both our Gram-positive and Gram-negative model bacteria showed that they could take up the miRNAs. The incorporation of eukaryote miRNAs in bacteria is consistent with their ability to absorb environmental nucleic acids, such as extracellular DNA, through natural competence. Alternatively, *in planta*, in the same way that some bacteria secrete small RNAs in outer membrane vesicles [31], bacteria may internalize external DNA *via* vesiduction [32], *i.e.* membrane fusion of a vesicle containing DNA or RNA. The use of vesicles seems, however, not necessary for plant-microbe miRNA-based interactions, as naked miRNAs were efficiently taken up by bacteria, and led to transcriptomic and community shifts.

In plants, miRNA induce mRNA cleavage or translation inhibition, through near perfect sequence complementarity [33]. Studies in the human gut also suggested that host miRNA interacts with bacterial mRNA through sequence complementarity [6, 8], so we used the rules for plant miRNA based on sequence homology, to search for targets in bacterial genomes. Using this model to select our target genes for the *in vitro* miPEP experiment, we found that two out of three targets were indeed inhibited following miPEP application and increased expression of the precursor miR159c. The third gene, encoding for alpha-2-macroglobulin, was, however, overexpressed in the presence of the miPEP. In bacteria, non-coding small RNAs can sometimes induce expression of target mRNAs [6, 31].

Even though the miPEP experiment showed that prediction tools worked well when focusing on a few genes, *Variovorax paradoxus* differentially expressed only 4 of the 137 predicted targets and differentially expressed another 247 non-predicted genes when exposed to a mixture of six rhizospheric miRNAs. This suggests that either 1) the bacteria adjusted their transcriptome in response to the shift in the expression of the miRNA-targeted gene, 2) as it is often the case in plants [34], the miRNAs targeted bacterial transcription factors, such as the ones from the LysR family that were differentially expressed at both time points, 3) the miRNAs led to the production of secondary siRNA, as previously shown in plants for two of our rhizospheric miRNAs, miR165 and miR161.1 [35], 4) target genes were translationally repressed, which would be undetectable with transcriptomics, though this is less common in plants than mRNA cleavage [33], 5) miRNAs protected targets from repression, as it was shown for arbuscular mycorrhizal fungi [36], or 6) since many predicted targets of the rhizospheric miRNAs were not in CDS, miRNAs could have affected DNA methylation [37] or interacted with gene promoters [38]. Clearly, further investigation is needed to clarify how eukaryotic miRNAs affect bacteria.

In contrast to *Variovorax*, plant miRNAs did not impact the transcriptome of *Bacillus*. In our simplified soil community, only 4 out of the 20 ASVs were significantly impacted by plant miRNAs, and, similarly, intestinal miRNAs only impacted the growth of specific bacterial strains [6]. Plant exosomes containing miRNAs are also preferentially taken up by some bacteria, affecting their gene expression and activity [8]. Our experiments were, however, carried out using naked miRNAs, excluding this explanation. We first thought that this could be related to differences in the cell wall that made miRNA entry impossible for Gram-positive, but our microscopy work disproved that. Another possibility is that the mechanism of interaction differs between the two groups of bacteria. In bacteria, chaperone proteins such as RNA-binding Hfq, ProQ, or CrsA proteins protect small RNAs and stabilize their interaction with mRNA, improving the formation of sRNA-mRNA duplexes that lead to gene silencing [39]. This crucial role of chaperones in sRNA-mediated interactions was only reported for a handful of Gram-positive bacteria [40, 41]. Alternatively, competence for DNA uptake depends on environmental conditions, such as stress, nutrient availability, and cell density [42]. The two bacteria tested might have different cues to initiate nucleic acid uptake, and the growing conditions might not have been met during the transcriptomic experiment to trigger this behavior in *Bacillus*. In any case, the differential transcriptomic response to miRNAs of the two bacteria tested suggests a selective mechanism in the rhizosphere. Any effect on a bacterium could have cascading effects on the rest of the community.

*Arabidopsis* miRNAs impacted the bacterial community in the root environment. We conducted three complementary experiments to prove this point. First, we examined the root-associated bacterial community of *A. thaliana* mutants affected in the biosynthesis of miRNA and/or siRNA. Many of these mutants had disrupted bacterial communities compared to wild-type plants. One of the most relevant mutants, the *dcl1-2* mutant, which is specifically impaired in miRNA production, was severely affected in its community composition at the phylum level and harbored a more diversified community. Microbial communities in the roots and rhizosphere of *ago1-27* and *RTL1myc* mutants resembled those of an unplanted soil more than those of wild-type plants. This suggests that mutations in small RNA related pathways lead to a certain dysbiosis in the roots and rhizosphere microbiota. Bacterial diversity was higher in the root environment of mutant plants which could reflect a weaker selection from these plants because of their lack of miRNAs and/or siRNAs. The mutations used are, however, pleiotropic, and plants were severely affected in their phenotype, which could have also affected the bacterial community.

Second, the bacterial community associated with soil-grown *Arabidopsis* responded significantly to miPEP application. As we showed in the *in vitro* experiment using miPEP159c, miPEPs stimulate the production their corresponding miRNA in plant tissues. This means that the up-regulation of a single miRNA could lead to changes in the bacterial community. Other than the direct effect of the overexpressed miRNA on the bacteria, the shifts observed in bacterial community could, however, also be explained by other factors. For instance, plant miRNAs alter various physiological processes within the plant, such as root development and plant immune response [34]. These changes induced by the overexpression of some miRNAs would also lead to large shifts in the plant microbiota.

Third, to exclude most of the indirect plant-mediated effects of the mutant and the miPEP experiments, we did an *in vitro* experiment with a simplified soil-derived bacterial community of twenty ASVs exposed to a mixture of synthetic miRNAs. The plant miRNAs affected the abundance of five ASVs and the growth of the community during the log phase. This shows that plant miRNAs directly affect bacterial communities. At the individual level, a bacterium could change in relative abundance because of 1) direct effect of the miRNA on its growth or 2) changes in the relative abundance of other bacteria with which it interacts. Some species can have a keystone role in interaction networks [43], and shift in these species would influence the entire community. For instance, the presence of *Variovorax* in the rhizosphere of *Arabidopsis* counteracted the root growth inhibition induced by many other members of the community [44]. Plant miPEPs, through their effect on miRNAs, also modulated the interactions between plants and key root symbionts, such as arbuscular mycorrhizal fungi [36] and rhizobia [45]. This effect would profoundly alter the microbial community, even if the miRNA had only affected a single keystone species.

We showed here for the first time that plant miRNAs are found in the rhizosphere of two model plants, that they are internalized in rhizosphere bacteria, that they affect the transcriptome of rhizosphere bacteria, and that they modulate soil microbial community composition and growth. The rhizosphere effect is thought to be mainly due to the rhizodeposition of various small organic molecules. Our study suggests a novel molecule, miRNAs, that plants could use to interact with their microbiota, challenging the current paradigm of rhizosphere microbial assembly. Many questions remain to be answered, but our study has shown beyond any doubt that plant miRNAs can shape the rhizosphere bacterial community.

## Acknowledgments

We want to thank Hervé Vaucheret and Taline Elmayan (IJPB, INRAE, Versailles, France) for sharing their *Arabidopsis* mutant seeds with us and Paul M. Orwin for providing the *Variovorax paradoxus* EPS strain. We also thank Christophe Penno for the helpful discussions. We also thank Jessy Tremblay (INRS, Laval, Canada) for the expertise and support regarding the confocal microscopy and flow cytometry experiments.

## Data availability statement

We deposited the sequence data generated in this study under NCBI BioProject accessions PRJNA836586 (rhizosphere miRNAs), PRJNA1107220 (root miRNAs), PRJNA1111839 (soil-extracted bacteria miRNAs), PRJNA1111831 (mutant and miPEP 16S rRNA gene amplicons), PRJNA1111829 (simplified soil community 16S rRNA gene amplicons), and PRJNA1111827 (*Bacillus* and *Variovorax* transcriptomics). The R code used to manipulate the data and generate the figures is available on GitHub (https://github.com/le-labo-yergeau/Middleton_miRNA) with the accompanying data being available on Zenodo (https://zenodo.org/doi/10.5281/zenodo.11105307).

---

**Figure S1.**
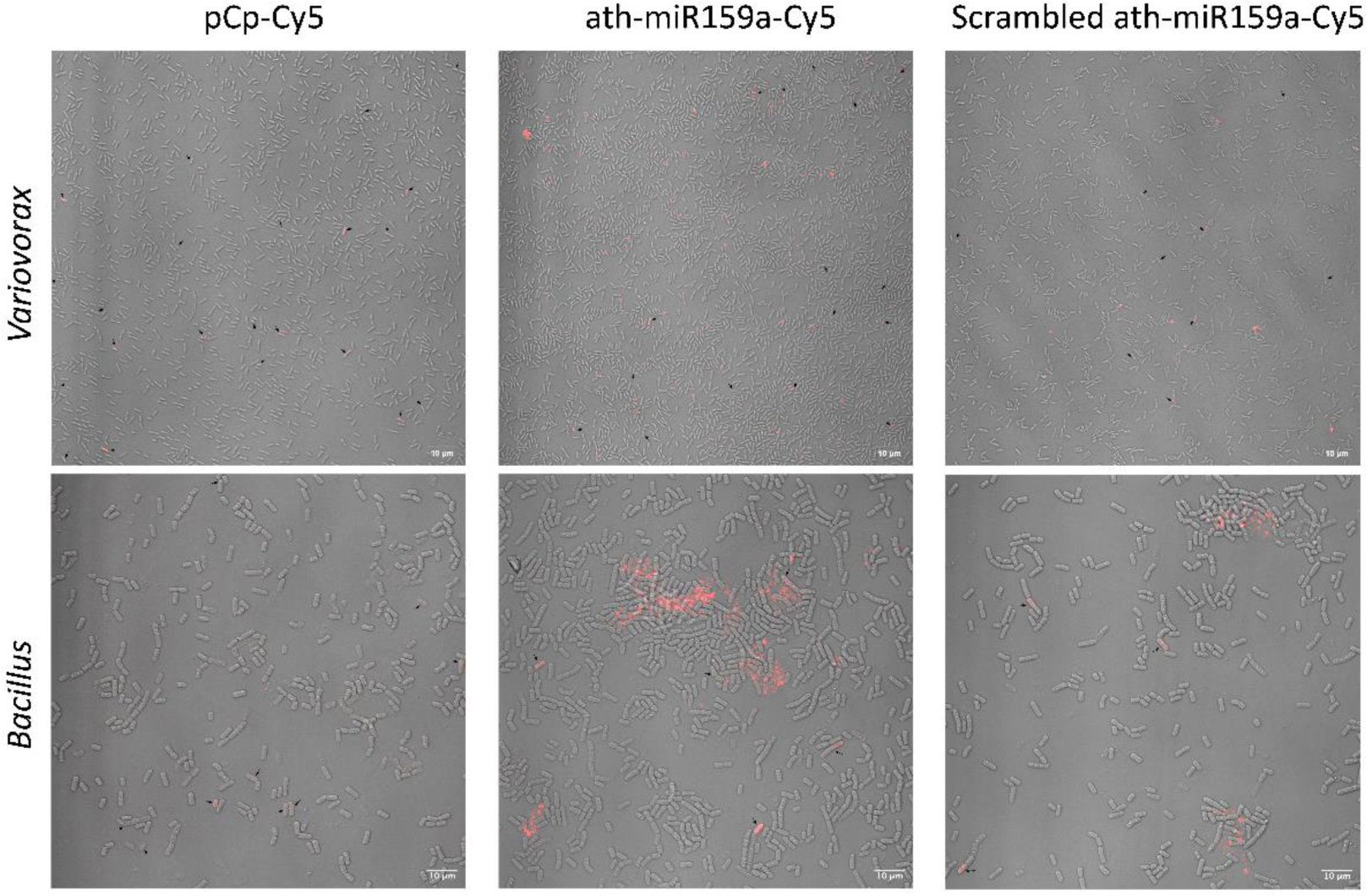

**Figure S2.**
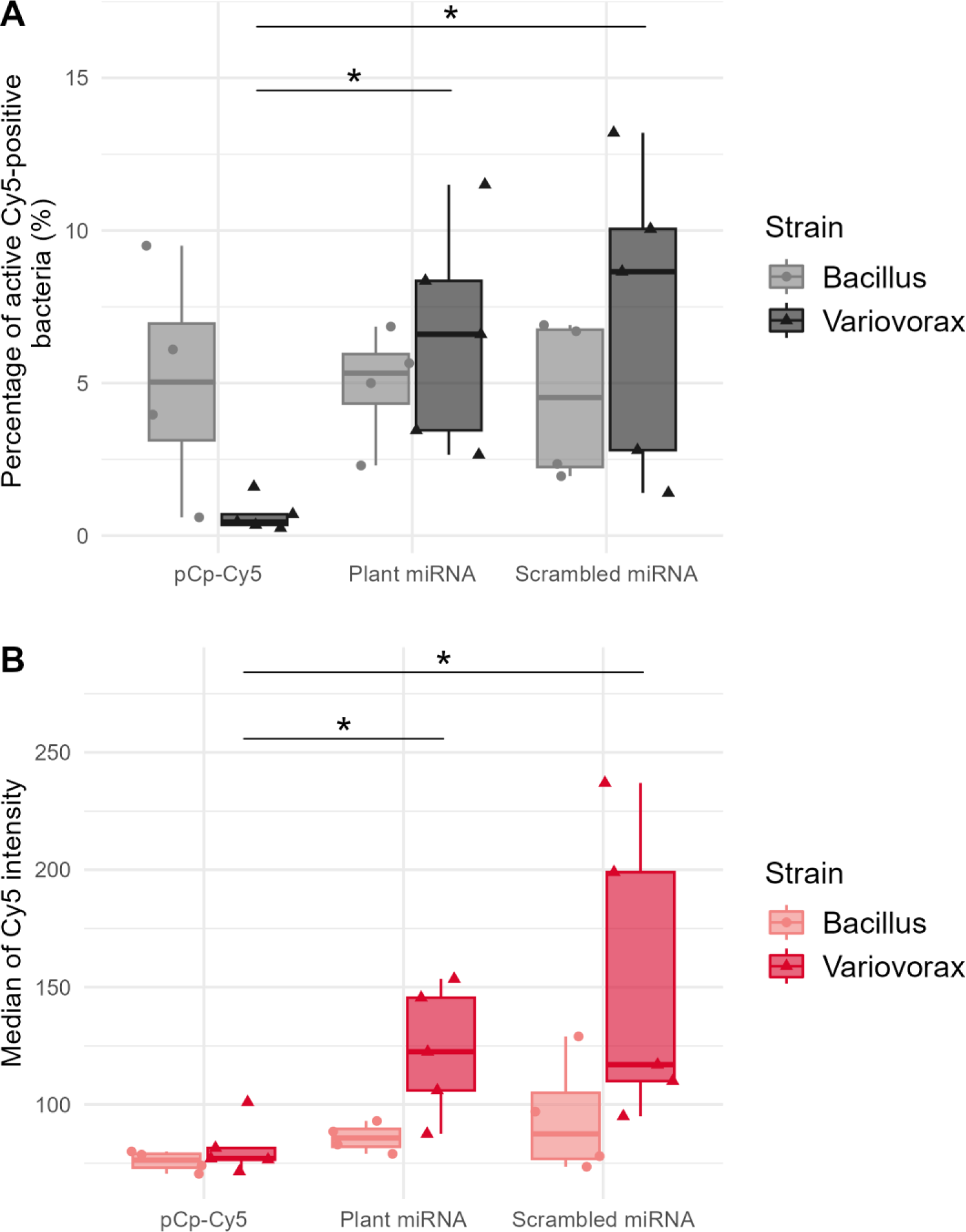

## Supplementary methods

### Detection of plant miRNAs in the roots and the rhizosphere

#### Plant growth conditions

For the detection of miRNA in the rhizosphere, *Arabidopsis thaliana* (Col-0) and *Brachypodium distachyon* (Bd21-3) were planted in triplicates and grown for a month in a growth cabinet, alongside unplanted soils. For the detection of miRNA in roots, *Arabidopsis thaliana* (Col-0) was planted in ten different experimental blocks and grown for 21 days in a growth cabinet, alongside unplanted soils. The program of the growth cabinets was: 12 hours of daylight, 3 hours of twilight, 3 hours of dawn light, and 6 hours of darkness at a constant humidity of 70%. The temperature varied in accordance with the light cycle between 20-25 °C.

#### Sampling, RNA extraction and sequencing

The rhizospheres and control soils were sampled and flash-frozen in liquid nitrogen. RNA was then extracted from 2 g of soil using MOBIO RNA Power soil kit, eluted in 100 µL, following the manufacturer’s guidelines. The roots were gently and quickly washed with sterile PBS, placed in a sterile tube and flash frozen with liquid nitrogen. The roots were lyophilized for three hours, crushed with a micro pestle, and RNA was extracted using the Qiagen RNeasy Plant Mini kit, following the manufacturer’s guidelines. The RNA was treated with DNase (Turbo DNA-free kit, Ambion), purified and concentrated (RNA Clean & Concentrator 25μl, ZymoResearch). We evaluated RNA quality with Nanodrop and 4% agarose gels, before sending it for sequencing on an Illumina HiSeq 2500 in single read mode (1×50bp) at the Centre d’expertise et de services Génome Québec (Montreal, Canada).

#### Bioinformatic analyses

Reads were trimmed using Trimmomatic [1] (v0.39) to remove the sequencing adapters with the following Illumina clip settings :5:10:4 and to remove the 5’ and 3’ flanking sequences containing bases having quality scores below 33 and 30, respectively. Common contaminants, such as Illumina adapters and PhiX spike-in sequences, were removed using BBDuk (BBMap v38.11, https://github.com/BioInfoTools/BBMap), with key parameters: k=20 and minkmerhits=1. Remaining sequences were then filtered by size such that reads having a length between 18 and 27 nucleotides were selected. Selected reads were mapped against either *A. thaliana* (TAIR10/GCA_000001735.1) or *B. distachyon* (bd21/GCF_000005505.1) reference genomes, which are the same as those used in miRbase [2], using BWA-aln [3] with the following parameters: mismatch=1 and seed=5. Using BEDTools v.2.29.2 [4] with -F 0.95 argument, a number of reads aligned inside miRNA coordinates were determined, meaning that at least 95% of each read had to belong inside the miRNA coordinates to be considered a valid hit. These assigned miRNAs which were mapped against the reference database were kept for abundance estimation across samples.

#### Statistical analyses

Rhizospheric and root miRNAs were defined as the miRNAs that had at least 10 reads for each the rhizosphere or root samples and a maximum of one read across all the bulk soil samples.

### Internalization of plant miRNAs by bacteria

#### Detection of plant miRNAs in rhizospheric bacteria

##### Plant growth and bacteria isolation

*A. thaliana* Col-0 was grown for a month, then the rhizosphere was sampled. Bacteria were isolated from 5 g of rhizospheric soil, using a density gradient. Samples were added to 30 mL of 0.2% sodium hexametaphosphate and rotated on a spinning wheel for 2 h. Samples were then centrifuged at 18 *g* for 1 min at 10°C, the supernatants were pooled two-by-two and centrifuged at 2824 x *g* for 20 min at 10°C. The microbe-containing pellet was resuspended in 10 mL of 0.8% NaCl and samples were again pooled by two. The resuspended solution was then transferred onto 10 mL of sterile Nycodenz and was centrifuged at 3220 *g* for 40 min. The bacterial layer was collected and a volume of 0.8% NaCl sufficient for a total of 35 mL was added. This solution was centrifuged at 3220 *g* for 15 min at 10°C. Finally, the pellet was resuspended in 1 mL of NaCl 0.8% to wash the residual Nycodenz away and centrifuged at 1700 *g* for 5 min at 10°C. The final pellet was resuspended in 100 µL TE buffer and kept at −20°C. Eight bacterial samples were obtained from the rhizosphere and six from unplanted soils.

##### RNA extraction and sequencing

RNA was extracted from this bacterial solution as follows: the 100 µL were transferred into a MP Biomedicals™ Lysing Matrix E tube with 250 µL of warmed 10% CTAB (Cetyl Trimethyl Ammonium Bromide) - 0.7 M NaCl, followed by 250 µL of 240 mM K_2_HPO_4_/KH_2_PO_4_ (pH 8.0) and 500 µL of phenol-chloroform-isoamyl alcohol (25:24:1) (pH 8.0). Samples were then bead-beaten using TissueLyser II (QIAGEN) at 30 Hz for 3 min. The tubes were centrifuged at 16000 *g* for 10 min at 4°C and the supernatant was transferred into a new tube. One volume of chloroform-isoamyl alcohol (24:1) was added to the supernatant and centrifuged again at 16,000 *g* for 10 min at 4°C. The supernatant was collected and 2 volumes of 30% PEG 6000-1.6 M NaCl were added, and the tube was inverted to mix and was left at 4°C for at least 2 h. The tubes were then centrifuged at 18,000 *g* for 30 min at 4°C to precipitate RNA. The RNA pellet was then washed with 500 µL of 70% ethanol and centrifuged at 18,000 *g* for 10 min at 4°C. The pellet was then air dried and resuspended in 50 µL of RNAse-free water. RNA quality and quantity was verified using a 1% agarose gel and Nanodrop, and then sent for sequencing on an Illumina NovaSeq 6000 using paired-end 100 bp with size-selected library.

##### Bioinformatic analyses

Sequencing results were processed as stated above with the rhizospheric miRNAs. To confirm that miRNAs found in the rhizospheric bacteria originated from the plant and not the bacteria, miRNA sequences were searched for on the + and - strands of 3,837 bacterial genomes [5], of which 1,160 bacteria were isolated from plants. These miRNA sequences were also searched for in the genome of *A. thaliana* TAIR10.1 in which they were found at their expected position.

##### Statistical analyses

We used the same strategy as above to define the internalized rhizospheric miRNAs, but with a lower threshold because there were less plant miRNA reads produced for this experiment. We defined the internalized miRNAs as the miRNAs that were represented by at least 5 reads in each rhizosphere microbial pellets and absent in the bulk soil microbial pellets.

#### Confocal microscopy

*Variovorax paradoxus* EPS and *Bacillus mycoides* YL123 were cultured on solid media (Bacto Yeast Extract Agar for *V. paradoxus* EPS and Tryptic Soy Agar for *B. mycoides* YL123) for 48 hours and then a few colonies were inoculated in 5 mL of the equivalent liquid media and grown overnight (250 rpm 28 °C). The cultures were then standardized to an OD600 of 0.200, treated with one of the 3’-Cy5-labelled oligos (plant ath-miR159a-Cy5, scrambled ath-miR159a-Cy5 or a modified version of cytidine pCp-Cy5, final concentration 2 µM) (Supplementary Table S1) and incubated for a total of 3 hours and 40 minutes (the cultures were within the log phase) after which they were treated with MitoTracker Green FM (Invitrogen, final concentration 2 μM). The cultures were immediately reincubated for 20 minutes, pelleted, washed with sterile PBS, pelleted again, and concentrated 10 times in sterile PBS. The cells (4.5 μl of washed and concentrated culture) were then placed on a glass bottom dish, covered with a thin slice of 2% agarose, and visualized using a confocal microscope (Zeiss LSM780). The experiment was repeated three independent times.

#### Flow cytometry

We prepared the bacterial cultures as for the confocal microscopy. After that, we took 5.5 μl of washed and concentrated culture and incubated it for 1 h and 30 min (room temperature). Then, we treated the bacteria with 4% Paraformaldehyde (PFA) for 40 min to fix them. We then stained the cells with a DNA marker (Hoechst 33342) for 35 min. After which we diluted the cells in sterile PBS to reach a final volume of 350 µl and processed the samples with a flow cytometer (BD LSRFortessa). For each sample, 50 000 events were measured. We used a variety of controls, such as cells only stained with Hoechst, cells only stained with MitoTracker GreenFM, cells stained with Hoechst + Mitotracker GreenFM and cells without dye to set appropriate gates and to ensure that our analyses were conducted on bacteria that were positive for our green (MitoTracker), red (Cy5 tagged miRNAs) and blue (Hoechst) markers. For each experiment, duplicates for each strain were prepared. The experiment was repeated five independent times for *V. paradoxus* EPS and four times for *B. mycoides* YL123. For our analyses, we used the mean of the duplicates and tested the differences between the strains and treatments with two Kruskal-Wallis tests. As a post-hoc test, we used Dunn’s test for pairwise multiple comparisons (significance=p adjusted < 0.05).

### Effect of miRNAs on bacterial gene expression

#### In silico analysis of miRNA targets

To predict potential plant miRNA targets in bacterial genomes, we implemented a workflow named *mirnatarget 1.0 : miRNA target finder* (https://github.com/jtremblay/MiRNATarget), which was inspired from the plant miRNA target finder, *psRNAtarget* [6]. This tool is based on specific pairing patterns between plant miRNA and targets [7] and implements SSEARCH36 (from the fasta36 v36.3.8 package) which uses the Smith-Waterman local alignment algorithm, resulting in optimal alignments [8]. Multiple targets for each miRNA can be identified. Based on the *psRNAtarget* rules for miRNA-target recognition, an *e*-value was calculated for each alignment and only those with an *e*-value ≤ 5 were retained. The positions of the targeted regions were noted with respect to the neighbouring coding sequences (CDS): the target sequence could be inside a CDS, at the 5’ or 3’ flanking regions (FLR) or in other short/very short inter-CDS regions, overlapping CDSs or elsewhere, further away from any CDS. Targets in the latter category were positioned at a distance superior to 100 nt from the 5’ end or at a distance superior to 350 nt from the 3’ end of the CDS. These distances were chosen as they correspond to the longest UTRs that are implicated in gene regulation. Targets in this category were discarded from the analysis, because it was considered that a miRNA in that area would not affect mRNA translation. The annotation of targeted regions and extraction of the closest CDSs were carried out with our tool “align2cdsRegions” (https://github.com/ntzv-git/align2cdsRegions). Sequences of targeted CDSs were retrieved using the *getfasta* tool from BEDTools [4] (v2.30.0) and GO (Gene Ontology) terms were attributed using InterProScan (v5.59-91.0) [9]. For each domain of GO (*i.e*. Biological Processes, Cellular Component, Molecular Function) and for each targeted CDS, a GO ancestor and their description were associated, and a level of hierarchical ontology was attributed [10].

#### Bacterial transcriptomic response to miRNAs

##### Bacterial growth and miRNA treatment

*Variovorax paradoxus* EPS was grown in 5% yeast extract (YE), at 30°C with 250 rpm shaking. *Bacillus mycoides* YL123 was grown in tryptic soy broth (TSB), at 25°C with 250 rpm shaking. Bacterial stocks were plated on YE or TSB and single colonies were selected for liquid culture overnight (*n* = 3), in 20 mL YE or TSB. In the morning, cultures were normalised as such: the optical density (OD) of overnight cultures was measured at 600 nm, which were then centrifuged at 5000 rpm for 5 min to collect the bacterial pellet. The supernatant was removed, and a calculated volume of medium was added to the pellet to reach OD = 1 for *B. mycoides* (*i.e.* 7.36 x 10^8^CFU/mL) and OD = 0.3 for *V. paradoxus* (*i.e.* 9.99 x 10^8^CFU/mL). Bacterial cultures were left to grow until the exponential phase was reached, which was determined previously (4-5 h for both strains). OD was then measured to check that the maximum number of cells/mL for RNA extraction was not exceeded (10^9^ cells/mL). At this point, 1 µg of the miRNA mix, or scrambled mix was applied to the cultures and gently mixed (15 µL of each miRNA solution diluted at 10 µM). Cultures were normalised to reach the same number of bacterial cells per mL. The volume of the cultures was varied to mimic natural differences in rhizosphere volume. Total concentration of miRNAs ranged between 65 nM-104 nM for *B. mycoides* and 45 nM-85 nM for *V. paradoxus*. The same cultures were sampled after 20 min and 120 min of incubation, to evaluate the evolution of gene expression in a same biological sample in response to a short and long confrontation with miRNAs. The miRNA mix was composed of 6 rhizospheric miRNAs : miR159a, miR159b, miR159c, miR161.1, miR158a and miR165b (Supplementary Table S1). The scrambled mix was composed of 6 RNA sequences with the same nucleic acids as the respective miRNAs but in a random order (Supplementary Table S1). To resemble mature plant miRNAs found in the rhizosphere, these synthetic miRNAs were single-stranded with a 3’ methyl group. They were synthesised by Integrated DNA Technologies (IDT).

##### RNA extraction

After incubation, 1 mL of every culture was sampled and centrifuged at 5000 rpm for 5 min. The bacterial pellet was resuspended in 100 µL of a lysozyme solution: 1 mg/mL for *V. paradoxus* (Gram −) and 3 mg/mL for *B. mycoides* (Gram +). After vortexing the resuspension, the samples were left to incubate for 15 min at room temperature. The following steps used solutions from the RNeasy Plant Mini Kit (QIAGEN). The guanidine-thiocyanate-based buffer RLT was added to the samples for further lysis (600 µL) and vortexed for 10 s. For *B. mycoides* samples, an additional step was necessary: the 700 µL were transferred to a sterile 2 mL Safe-Lock tube containing 100 mg glass beads (0.1 mm diameter) and 100 mg zirconia/silica beads (0.5 mm diameter). Beads were resuspended by pipetting then cells were disrupted using Fast Prep for 50 s (5 m/s), followed by 5 min on ice and another 50 s of Fast Prep. Tubes were centrifuged at 10 000 *g* for 1 min and supernatant was transferred to a new tube. The next steps were common to both strains. An equal volume of 70% ethanol was added to the lysed samples and mixed by pipetting. The lysate was transferred to a RNeasy spin column (pink column) in a 2 mL collection tube and centrifuged at 10 000 rpm for 30 s, then the flow-through was discarded. Columns were first washed with 700 µL of the RW1 wash buffer, followed by 500 µL RPE buffer and again with 500 µL RPE buffer. Collection tube was changed, and the column was centrifuged for 1 min at maximum speed to dry the membrane off. To elute RNA, columns were placed in new 1.5 mL tubes and 30 µL of RNAse-free water was added before centrifugation. The 30 µL were then reloaded and centrifuged one last time.

##### RNA-seq & sequence analysis

Total RNA was sent for sequencing at the Centre d’expertise et de services Génome Québec (Montreal, Canada), where ribosomal RNAs were depleted and messenger RNAs were sequenced. RNA-seq raw reads were processed with ShotgunMG [11]. Reads were trimmed and filtered for quality (Trimmomatic v.39) and mapped (BWA v0.7.17) against their respective reference genome. The genome of *B. mycoides* YL123 originated from our lab, was sequenced, processed and submitted to the NCBI through the Prokaryotic Genome Annotation Pipeline (PGAP) under the assembly accession GCA_024297165.1 on GenBank. The *V. paradoxus* EPS genome was published with accession number NC_014931 [12]. BedTools v2.23.0 was used to generate read count files from each .bam file that were then merged to generate a raw count matrix. The latter file was used as input data for a differential gene expression analysis using DESEq2 (Bioconductor, version 1.36.0). For each strain and each time point, gene expression of samples treated with synthetic miRNAs was compared to those treated with scrambled synthetic miRNAs.

#### In vitro miPEP transcriptomic experiment

##### Arabidopsis culture in vitro

Approximately 50 *Arabidopsis thaliana* col-0 seeds were surface-sterilised by soaking them in 70% ethanol for 5 minutes, followed by 20 minutes in 0.5% sodium hypochlorite. Seeds were then rinsed six times with sterile distilled water and resuspended in 1 mL of water, which was poured onto a Petri dish, with some culture medium, to let the water evaporate under sterile conditions. The Petri dish was sealed with parafilm and placed at 4°C for 3 days, for the stratification process, allowing a synchronous germination. The culture medium was composed of 1.2 g of Hoagland’s No. 2 basal salt mixture (Sigma-Aldrich, reference H2395), 8 g of agar ‘suitable for plant cell culture’ (Sigma-Aldrich, ref. A8678) and 500 mL of MilliQ water. The pH of the medium was adjusted with HCl to 5.8-5.9, before adding the final 250 mL of water, resulting in 750 mL of medium. The solution was autoclaved and poured into square petri dishes for plant culture. When hardened, around 2 cm of solid medium was cut out, creating a ridge where the surface-sterilized seeds were positioned (10-15 seeds/dish). Petri dishes with these positioned seeds were sealed with microtape and placed vertically in a growth chamber, with 16 h of daylight at 21°C, 8 h of darkness at 19°C, with 25% relative humidity.

##### miPEP treatment

Twenty days after the seeds were placed in the growth chamber, the seedlings had developed at least 6 leaves and were ready to be treated. The miPEP treatment consisted in applying 100 µL of a 10 µM solution to each plant, starting at the tip of the root to the base (bottom-to-top). A second treatment was applied 24 h later, before the microbial inoculation. There were three conditions: miPEP159c treatment (MQNLRVHVFLIESARC), a scrambled version of miPEP159c and a water control, each treatment was applied to 7 Petri dishes (biological replicates), containing 10-15 plants each (pseudo-replicates). The use of a scrambled control as well as a solvent control (*i.e.* water) was to ensure that any “miPEP-effect” was indeed due to its action on the plant and not *via* some nutritive or anti-microbial effect of the peptide.

##### Bacterial growth and inoculation

*Variovorax paradoxus* EPS was first inoculated, from a −20°C stock, on a yeast extract Petri dish (5 g/L) to check that colonies are homogenous. A single colony was transferred into a liquid culture of yeast extract to grow overnight at 30°C and 110 rpm. The overnight culture’s optical density was measured at 600 nm to estimate the number of bacterial cells added to each plant: at OD = 1, the estimated cellular concentration is 3.33 x 10^9^ CFU/mL. 100 µL (with OD = 0.418 *i.e.* ∼1.39 x 10^8^ cells) of the overnight culture was directly applied to each plant, from the tip to the base of the root, one hour after the second miPEP treatment.

##### Sampling

Two hours after the bacterial inoculation, the plants and rhizospheres were sampled. The rhizosphere samples were isolated using a sterile spatula by extracting small squares of medium around the tip of the roots. Whole plants were gently extracted from the medium, by pulling on the aerial system. Samples were flash-frozen in liquid nitrogen and ground shortly after, using a sterile mortar and pestle. The resulting fine powder was stored at −80°C.

##### RNA extraction

The NucleoSpin ® RNA kit (MACHEREY-NAGEL, Düren, Germany) was used to extract RNA from ∼100 mg of previously ground powder, with some changes to the manufacturer’s protocol. During the cell lysis, 400 µL of RA1 solution were used instead of 350µL, followed by 400 µL of ethanol to adjust the RNA binding conditions. The following steps remained the same, up until the elution, where only 30µL of water was used and left to incubate on the column before centrifugation, resulting in better yields. DNase reaction was performed on-column during the protocol.

##### Target prediction

Potential targets of miRNA159c in the genome of *Variovorax paradoxus* EPS were identified using the online *<u>psRNAtarget</u>* tool. On the server, the ath-miRNA159c sequence (UUUGGAUUGAAGGGAGCUCCU), obtained from <u>miRBase</u>, and the accession number of the EPS genome (CP002417.1) were uploaded for analysis, with the default parameters of the Schema V2 of the 2017 release. Three of the targeted genes were selected for *in vivo* quantification.

##### RT-qPCR

Retrotranscription was performed using 50 ng of total RNA, more or less diluted to start the reaction with 12 µL of RNA, to which was added 1 µL dNTPs (10 mM) and 1 µL of random primers (200 ng), followed by a 5-minute incubation at 65°C. Samples were then placed on ice for 1min. Sequentially, 4 µL of 5X First-Strand Buffer, 1 µL of DTT (0,1 M) and 1 µL of SuperScript™ Reverse Transcriptase (200 U/µL, Invitrogen) were mixed in by pipetting and the samples incubated for 5 min at 25°C, followed by 1 h at 60°C and 15 min at 70°C. Samples were then placed on ice and 100 µL of RNAse-free water was added to the 25 µL of newly synthesised cDNA. Quantitative PCR was performed using a reaction mix composed of 5 µL of iTaq Universal SYBR Green Supermix (Bio-Rad Laboratories, ref. 1725124), 0.6 µL of primer mix (10 µM, forward & reverse) and 3.4 µL of RNAse-free water, to which 1 µL of cDNA was added. All measurements were performed using three technical replicates, in 384-well plates, on a Roche LightCycler ® 340 thermocycler. The qPCR was programmed as follows : a first step of polymerase activation and DNA denaturation at 95°C for 5 min; then the cycle commenced with further denaturation at 95°C for 10 sec, an annealing step at Tm for 20 sec and an extension step at 72°C for 30 sec; after 40 cycles, a final melting curve was produced: from 65°C to 97°C, with 5 acquisitions per °C. The melting temperature (Tm) for each primer set was adjusted to optimise the amplification specificity and their efficiency was validated beforehand and can be found with primer sequences in Supplementary Table S2. Using a dilution range of a cDNA mixture, each primer set’s efficiency was calculated using the slope of the *x*=log(dilution); *y*=Ct, in the following equation: (10^(−1/(slope))-1)*100, which should be in the 90-110% range. A melting curve was produced at the end of each qPCR program to ensure that a single gene was amplified. In some cases, the amplicon was verified by gel electrophoresis.

##### Data analysis

After RNA extraction and RT, two samples were discarded from further analysis due to experimental mishaps, resulting in 6 replicates in the “miPEP” and “water” conditions and 7 replicates in the “scrambled” condition. Quantification of pri-miR159c only succeeded in 6 out of 7 “scrambled” replicates and 5 out of 6 “water” replicates. Bacterial genes, *i.e*. Lys R, phosphatidate and alpha-macroglobulin, were quantified relative to RecA and GyrA reference genes, whereas the plant pri-miRNA159c was quantified relative to a plant reference gene, cyclophilin. The number of PCR cycles necessary for the fluorescence to emerge from the background noise is referred to as the “*Ct*”. Noise was automatically determined by the LightCycler 480 software and *Ct*s were calculated with the Abs Quant/2nd Derivative Max - high confidence mode. A coefficient of variation (CV) was calculated for the three technical replicates and a threshold of 2% was applied : extreme technical replicates were excluded, if this limit was exceeded. The remaining technical replicates were averaged to create the *Ct* value for each sample. The analysis of pri-miR159c expression was done using the Pfaffl method, which takes into account the efficiency of each primer set [13]. This method calculates a relative expression ratio based on the difference of *Ct*s of a gene of interest in a sample *versus* a calibrator, and in comparison, with a reference gene. The analysis of bacterial genes relied on two reference genes, which called for a slightly modified version of the Pfaffl method. To calculate a relative expression ratio, we had to normalise our qPCR data using the geometric mean of both reference genes [14]. In both cases, we averaged the samples treated with water to create a calibrator.

##### Statistical analyses

Statistical analyses were performed on the resulting relative expression ratios, comparing group-to-group, using a Welch Two Sample t-test or a Wilcoxon rank sum exact test, depending on if the data were normal.

### Effect of miRNAs on the bacterial community

#### Arabidopsis mutant experiment

##### Plant growth and mutant description

Five *A. thaliana* mutants were grown in a mix of soil and sand (2:1) for a month, in individual pots. Mutants were provided by Hervé Vaucheret and Taline Elmayan (IJPB, INRAE, Versailles, France). *RTL1* mutant over-expresses RTL1 protein which results in a suppression of siRNA pathway without affecting miRNAs [15]. *RTL1myc* over-expresses RTL1 protein flagged with Myc epitope, rendering RTL1 less active, so siRNA pathway is less suppressed than with *RTL1* mutant. *Ago1-27* mutant has AGO protein function partially impaired and is completely post-transcription gene silencing (PTGS) deficient [16]. *Dcl1-2* mutant has total loss of function of DCL1 protein resulting in low levels of miRNA and developmental problems [17]. *Hen1-4* mutant is miRNA defective but is also affected in some siRNA – PTGS [18]. HEN1 methylates siRNA and miRNA to maintain their levels and size, but also to protect them from uridylation and subsequent degradation [19].

##### DNA extraction, sequencing and qPCR

We grew *A. thaliana* mutants and wild-type plants in quadruplicate for a month and sampled the root, rhizoplane and closely adhering rhizosphere. Two of the wild-type plants were lost during the experiment and subsequent analyses. DNA was extracted using NucleoSpin Plant II kit (Macherey-Nagel). DNA was sent for 16S rRNA gene amplicon (primers 341F: 5’- CCTACGGGNGGCWGCAG-3’ and 534R: 5’- ATTACCGCGGCTGCTGGCA – 3’) sequencing on an Illumina MiSeq in paired-end mode (2×250 bp) at the Centre d’expertise et de services Génome Québec (Montreal, Canada). In parallel, 16S rRNA gene was quantified by qPCR, using the same primers as for sequencing. Within each well was 0.1 µL of each primer (10 µM), 4 µL LightCycler® 480 SYBR Green I Master (final volume 6 µL PCR mix) and 2 µL of DNA (25 ng.µL^−1^). We used the following 40-cycle program: pre-incubation (95°C-4 min) / amplification (95°C – 30 s; 49°C – 1 min; 72°C – 1 min) / 72°C-10 min/ melting curve (95°C – 5 s; 49°C – 1 min; 97°C – continuous 5 measures/ °C) / cooling (40°C – 30 s). At the last step of amplification, a single measurement was performed and then continuously during the melting curve step. A minimum of 3 technical replicates was quantified. The number of copies of 16S rRNA gene was determined in comparison with a standard curve using serial dilutions of plasmids with cloned fragments (R² = 0.994).

##### Amplicon sequences processing

Amplicon sequencing data was processed with AmpliconTagger [20]. Briefly, contaminants and unpaired reads were removed, and remaining sequences were trimmed to remove adaptors and primers. Reads were then filtered for quality control such that reads having at least one N or having average phred score quality less than 20 were left out. A total of 3,776,292 reads passed the quality control and 1,230,944 sequences were successfully processed to generate ASVs (DADA2) [21]. ASVs were filtered for chimaeras using DADA2’s internal removeBimeraDeNovo (method = ‘consensus’) workflow followed by VSEARCH’s [22] UCHIME *de novo*. Each remaining ASV was assigned a taxonomic lineage by using the RDP classifier with the SILVA [23] R138. ASVs assigned to bacterial/archaeal and rendered into ASV abundance tables.

##### Microbial community analyses

Analyses of ASVs composing the microbial communities were performed using the R package “phyloseq” v 1.32.0 [24]. After checking the rarefaction curves, all samples seemed sufficiently sequenced. For Shannon diversity index calculation, all reads were rarefied to the lowest number of reads found in a sample. Significance of differences in alpha-diversity between mutants was tested using a linear model, whilst checking the normal distribution of residuals and using wild-type (WT) samples as a reference. If residuals did not follow a normal distribution, a general linear model (GLM) with gamma distribution and “log” or “link” scale were used. All tests performed were, by default, two-sided.

##### Statistical analyses

The effect of miRNA/siRNA mutation in *A. thaliana* on the structure of microbial communities was visualized using Principal Coordinate Analysis (PCoA) ordinations of Bray-Curtis dissimilarities. Permutational Multivariate Analysis of Variance (Permanova) was used to determine the statistical impact of miRNA/siRNA mutations on the microbiota structure, alongside checking group dispersion and pairwise tests.

#### miPEP experiment

##### Plant growth and miPEP treatment

*Arabidopsis thaliana* Col-0 were sowed on a mix of potting soil and sifted sand (<2.2 mm) (ratio 2:1), in a greenhouse. After germination, seedlings were transferred into individual pots (3 cm diameter, 5 cm depth) that were moved to a growth chamber under 16 h of daylight, 8 h of darkness, at 20°C. When the plants reached the 6-leaf stage, the miPEP treatments started. Plants (16 replicates/condition) were treated with 500 µL of water (control condition) or a miPEP solution (20 µM of miPEP159a: MTWPLLSLSFLLSKYV, miPEP159b: MGLRKVLEMNTIFDSLFLSH or miPEP159c: MQNLRVHVFLIESARC), applied at the base of the crown, 3 times a week for a total of 10 applications. After each application, the trays in the chamber were moved around to diminish any border effect.

##### Sampling & DNA-RNA extraction

Once the treatments were over, each individual pot was spilled on a sieve and the aerial part was separated from the root system. The roots of two plants were pooled together and briefly rinsed in 10 mL of PBS (1X). The roots and the attached soil were then separated from the rinsing solution, using a funnel and a sterile compress, and were subsequently dried using absorbent paper. The dried roots and rhizosphere were flash frozen in liquid nitrogen and stored at −80°C. DNA was extracted using the NucleoSpin® Plant II mini kit (MACHEREY-NAGEL, Düren, Germany) whereas RNA was isolated with a homemade protocol [25]. The integrity and quantity of the extracted DNA and RNA was estimated using Nanodrop and by running a 1% agarose gel. Amplicon sequencing, processing, qPCR quantification, and analyses were performed as described above.

#### Simplified soil community experiment

##### Soil microbes enrichment

The soil microbes were enriched from five different media: 1) Tryptic Soy Broth, 2) Potato Dextrose Broth, 3) Minimal medium + Carbon solution + NH4NO3, 4) Minimal medium + Carbon solution + urea and 5) Minimal medium + Carbon solution + amino acids. The minimal media were supplemented with a solution of artificial carbon-rich root exudates: 20 mM glucose 20 mM fructose, 12 mM sucrose, 20 mM lactic acid, 12 mM citric acid, 16 mM succinic acid. For these media, the final concentration was 0.5 g/L of C and 0.1 g/L of N. To culture the soil microbes, 2 g of sieved agricultural soil (sampled at the Armand-Frappier experimental field: 45.5416N, −73.7173E) was incubated in 20 mL of each media, for 28 h (200 rpm, 25 °C). The cultures were, filtered 30 µm, normalized to the same optical density, pelleted (4 °C, 15 min, 4700 *g*), suspended in PBS, pooled and aliquoted into sterile cryotubes. A cryoprotective solution 2x (0.6% (w/v) Tryptic Soy Broth, 10% (v/v) DMSO and 2% (w/v) trehalose) was added to the cultures (1:1 (v/v)) and the cryotubes were gently mixed (5x inverting), left to equilibrate for 20 min and placed at −80 °C [26]. Cells were revived, inoculated in 5 mL of TSB overnight, washed, normalized to OD600 0.200 and inoculated in a 96-wells plate containing a mixture of 17 amino acids as nitrogen source and artificial root exudates as a carbon source.

##### Microbial growth and miRNA exposure

To investigate the potential modification of microbial activity by plant miRNAs, we revived, cultured overnight (in 5 mL of TSB, 28 °C, 200 rpm), washed (2x) and normalized the soil microbes (OD600 0.200). We then cultured them in a medium containing an equimolar mixture of 17 L-amino acids (15 mM), artificial root exudates (15 mM) and a miRNA treatment that had a final concentration of 10 µM (five biological replicates were prepared). The miRNA treatment consisted of an equimolar mix of five miRNAs (ath-miR158a-3p, ath-miR158b, ath-miR159a, ath-miR827, and ath-miR5642b) of either the plant mimics or scrambled controls (Supplementary Table S1). Microbial activity was quantified by introducing a tetrazolium dye, which undergoes a color change to purple in the presence of dehydrogenases, subsequently intensifying the optical density at 600 nm. Optical density measurements were recorded hourly using a plate reader. The experiment was concluded after 52 hours.

##### 16S rRNA gene amplicon sequencing

To determine whether alterations in microbial activity were indicative of changes within the bacterial community, we proceeded with a microvolume physical DNA extraction [27] from the cultures and prepared the libraries to sequence the V4-V5 region of the 16S rRNA gene (primers 515F-Y: GTGYCAGCMGCCGCGGTAA and 926R: CCGYCAATTYMTTTRAGTTT) [28] using the Illumina MiSeq platform at the Centre d’expertise et de services de Génome Québec, Montreal, Canada. The sequences were processed with the AmpliconTagger pipeline [20], and analysed as described above.

---

**Supplementary Table S1:**
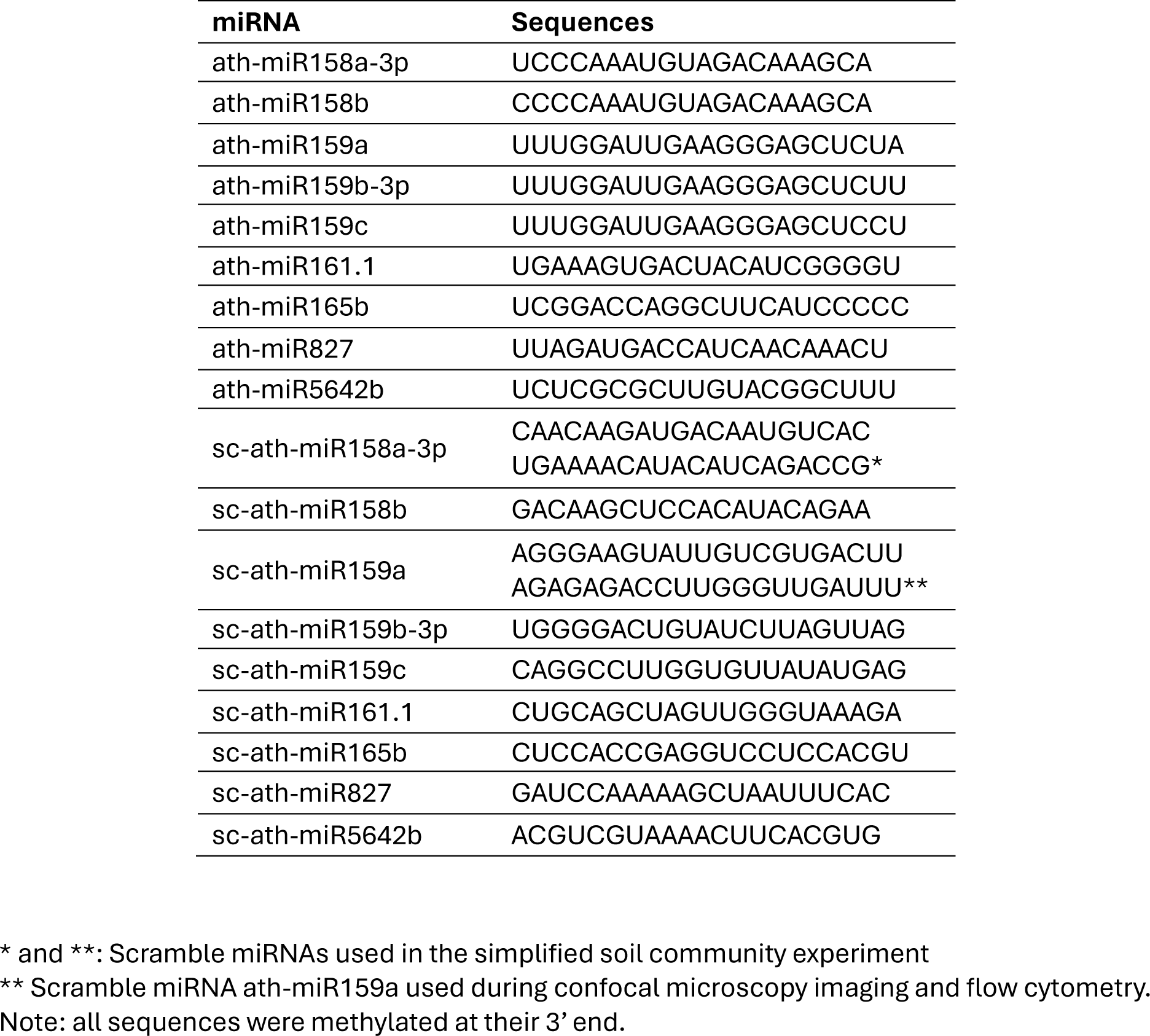
Sequences of the single-stranded synthetic miRNAs.

**Supplementary Table S2:**
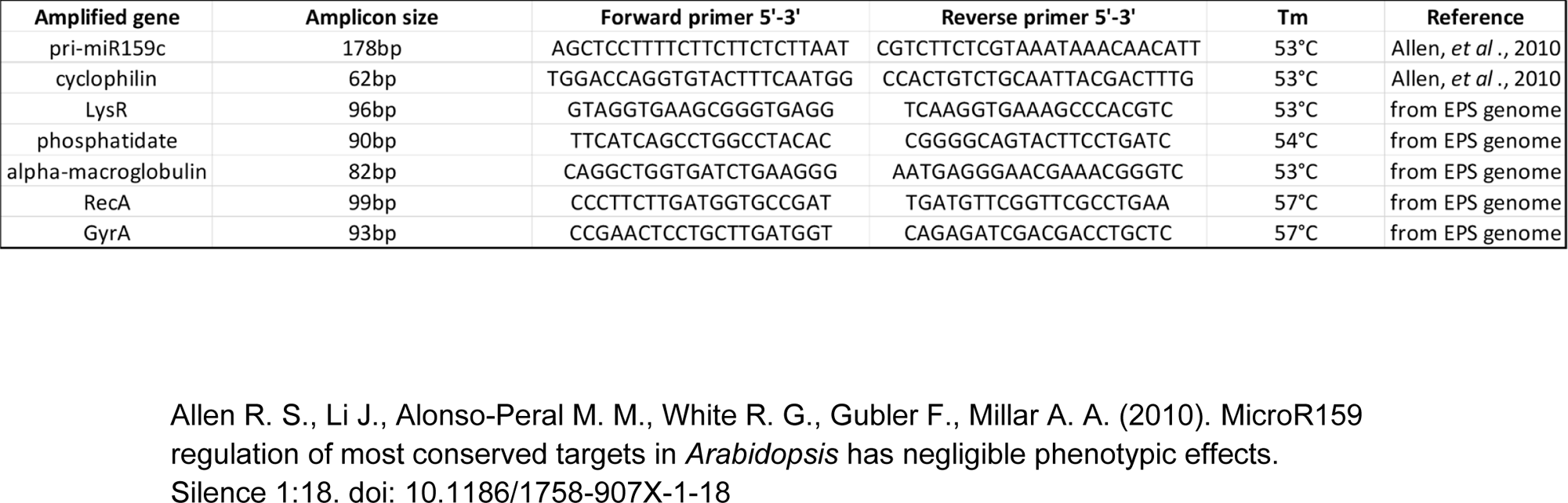
Primers used for qPCR in *Arabidopsis thaliana* and *Variovorax paradoxus* EPS for the *in vitro* miPEP transcriptomic experiment. Most primers were designed based on the EPS genome, except for plant gene primers.

